# Discovery of a CHI3L1-Targeted Small Molecule Modulating Neuroinflammation in Alzheimer’s Disease via DNA-Encoded Library (DEL) Screening^†^

**DOI:** 10.1101/2025.10.20.683491

**Authors:** Baljit Kaur, Longfei Zhang, Hossam Nada, Laura Calvo-Barreiro, Moustafa T. Gabr

## Abstract

Chitinase-3-like protein 1 (CHI3L1, also known as YKL-40) has emerged as a central effector of astrocyte-mediated neuroinflammation and a promising biomarker for Alzheimer’s disease (AD). However, small molecule CHI3L1 inhibitors that modulate neuroinflammation are limited. Here, we report the discovery of a CHI3L1-targeted small molecule, **DEL-C1**, identified through DNA-encoded library (DEL) screening and validated using orthogonal biophysical, computational, and cellular approaches. **DEL-C1** demonstrated direct CHI3L1 binding in microscale thermophoresis (MST) and surface plasmon resonance (SPR) assays, with reversible and concentration-dependent association. Molecular docking and 100-ns molecular dynamics simulations revealed a stable binding mode within the CHI3L1 substrate groove, anchored by Tyr206 and flanked by Trp99 and Trp352, supporting a thermodynamically favorable interaction. In vitro ADME profiling indicated a balanced physicochemical profile, permeability, and metabolic stability, consistent with CNS drug-like properties. Functionally, **DEL-C1** reversed CHI3L1-induced astrocyte dysfunction by restoring Aβ uptake, lysosomal acidification, and proteolytic activity, while reducing CHI3L1 and IL-6 secretion. **DEL-C1** also suppressed CHI3L1-driven NF-κB transcriptional activation, highlighting its anti-inflammatory potential. Collectively, this study establishes **DEL-C1** as a promising small molecule modulator of CHI3L1 and a chemical tool to interrogate astrocyte-driven neuroinflammation in AD.

## Introduction

Neuroinflammation is now recognized as a central driver of Alzheimer’s disease (AD) pathogenesis, operating alongside amyloid and tau pathology to shape disease progression and cognitive decline.^1-3^ The chronic activation of astrocytes and microglia creates a self-reinforcing inflammatory loop that exacerbates neuronal injury, synaptic loss, and impaired neurogenesis.^4,5^ Despite extensive evidence linking glial activation to neurodegeneration, current therapeutic strategies targeting inflammation, such as broad immunosuppressive drugs or cytokine inhibitors, have shown limited efficacy in clinical trials.^6,7^ These failures underscore the need for selective modulators that precisely reprogram pathogenic glial signaling without compromising essential neuroimmune homeostasis.

Among the molecules implicated in astrocyte-mediated neuroinflammation, chitinase-3-like protein 1 (CHI3L1, also known as YKL-40) has emerged as a key biomarker and potential effector of disease progression.^8-10^ CHI3L1 is a secreted glycoprotein produced predominantly by reactive astrocytes, and its levels in cerebrospinal fluid (CSF) and plasma correlate with tau pathology, cortical atrophy, and cognitive decline in patients with AD.^11-13^ Mechanistic studies have revealed that CHI3L1 is not merely a bystander biomarker but an active signaling mediator that modulates both astrocytic and neuronal function through distinct receptor pathways.^14-18^ Connolly et al. demonstrated that CHI3L1 is markedly up-regulated in astrocytes surrounding amyloid plaques and tau-positive regions, and that elevated cerebrospinal-fluid CHI3L1 levels correlate with cortical atrophy, tau pathology, and cognitive decline in individuals with AD.^14,18^ Building on these clinical and histopathological observations, Zeng et al. showed that astrocyte-specific deletion of CHI3L1 in 5xFAD mice significantly reduced amyloid-β accumulation, microgliosis, and synaptic loss, while rescuing learning and memory performance.^15^ Together, these findings establish CHI3L1 as a causal effector linking astrocytic reactivity to neurodegenerative progression.

Mechanistic insights from recent experimental models have revealed that CHI3L1 acts through dual receptor-mediated pathways that converge on neuronal and neurogenic dysfunction. Yang et al. reported that CHI3L1 secreted by reactive astrocytes binds to the chemoattractant receptor-homologous molecule CRTH2, activating the IKKβ–S6K1–S6 axis, which suppresses hippocampal neural-stem-cell proliferation and differentiation.^16^ In parallel, Song et al. demonstrated that CHI3L1 engages the receptor for advanced glycation end products (RAGE) in neurons, leading to GSK3β-mediated β-catenin destabilization, dendritic simplification, and synaptic loss.^17^ Pharmacological inhibition or genetic blockade of CHI3L1, CRTH2, or RAGE restored β-catenin signaling, neurogenesis, and cognitive performance in AD models. Collectively, these data position CHI3L1 as a central node in astrocyte-driven neuroinflammation, acting through distinct CRTH2- and RAGE-dependent pathways that represent promising therapeutic targets for AD.

Despite strong genetic and pathological evidence implicating CHI3L1 in AD, the potential of CHI3L1 as a druggable target has not been realized. Monoclonal antibodies and antisense approaches are limited by poor blood–brain barrier (BBB) penetration and high manufacturing complexity, while global inhibition of glial activation risks unwanted immunosuppression.^19,20^ By comparison, small molecules present several advantages, including efficient BBB penetration, suitability for oral administration, reduced immunogenic potential, and facile optimization of pharmacokinetic (PK) properties, which permits flexible dosing to minimize adverse effects.^21,22^ Notably, small molecule modulators capable of direct CHI3L1 engagement are limited. In this context, our previously reported CHI3L1-targeted small molecules had no impact on modulating neuroinflammation.^23-25^ These challenges highlight the urgent need for tractable chemical tools that can modulate CHI3L1 signaling in a targeted, reversible, and CNS-permeable manner.

DNA-Encoded Library (DEL) technology offers a transformative platform to identify such molecules by enabling affinity-based screening of billions of compounds against recombinant targets using minimal material input.^26-28^ DEL screening is particularly well-suited for glycoproteins like CHI3L1, whose shallow binding surfaces and dynamic conformations often preclude high-throughput screening (HTS) success. Our laboratory previously employed this platform to discover first-in-class small-molecule binders of SLIT2, validating DEL as a powerful approach for challenging extracellular protein–protein interactions.^29^ Extending this strategy to CHI3L1 provides an opportunity to uncover small molecules that directly bind and modulate this astrocytic effector of neuroinflammation.

Here, we report the discovery of a CHI3L1-targeted small molecule identified through DEL screening and validated using orthogonal biophysical, cellular, and computational approaches (Figure 1). We confirmed CHI3L1 engagement using and microscale thermophoresis (MST) and surface plasmon resonance (SPR) assays, established its functional activity using in vitro models of astrocyte-mediated neuroinflammation, and employed computational modeling to map its CHI3L1 binding interface. Finally, our workflow (Figure 1) involved preliminary evaluation of the in vitro PK profile of the CHI3L1-targeted hit to assess druggability and CNS suitability. This integrative workflow (Figure 1) demonstrates a scalable strategy for discovering first-in-class small molecule modulators of neuroimmune targets and establishes a framework for developing CHI3L1 inhibitors as potential therapeutics for AD.

**Figure 1.**
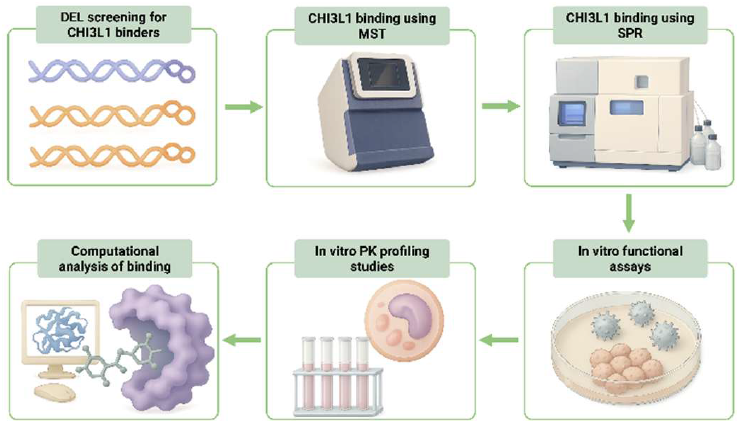
Schematic workflow for the identification of small molecule CHI3L1 binders as potential therapeutics for AD. Our workflow integrates DEL screening with orthogonal biophysical, cellular, and computational approaches.

## Results and discussion

### DEL screening

The DELopen platform developed by WuXi AppTec provides an open-access DEL system, allowing academic researchers to conduct affinity-based selections across a broad range of chemical diversity without proprietary restrictions. The platform encompasses multiple structurally varied libraries that collectively explore diverse regions of chemical space, encompassing numerous pharmacophores and scaffolds. A DEL screening campaign using the DELopen library (4.2 billion compounds) was conducted against the human CHI3L1 protein. The selection workflow of hit compounds involved exclusion of non-target control (NTC) signals (C4) from the target conditions (C1-C3), application of an enrichment score cutoff of 100, and prioritization of compounds that demonstrated exclusive binding to the CHI3L1 active region (A area) (Figure 2A).

**Figure 2.**
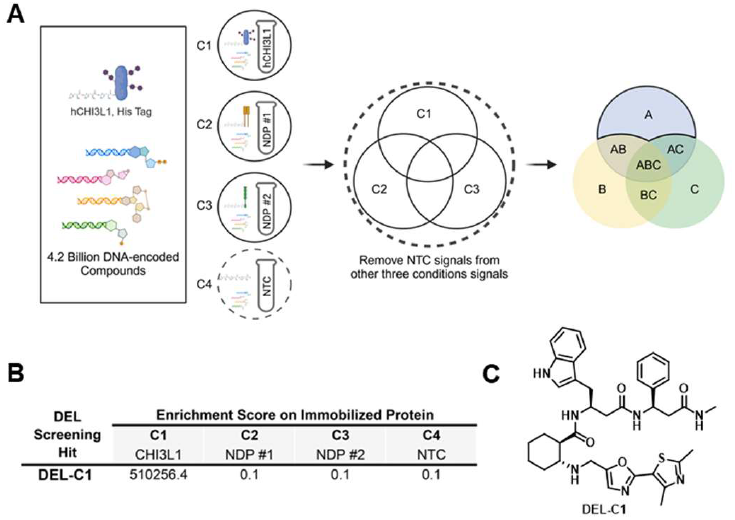
Discovery of lead CHI3L1-targeted small molecules DEL screening. **A**. Twenty-seven DNA-encoded libraries comprising approximately 4.2 billion compounds were screened against recombinant human CHI3L1 protein immobilized on HisPur™ Ni-NTA magnetic beads (C1). Two additional immobilized human proteins served as non-disclosed comparators (C2, C3), and the affinity matrix alone functioned as a non-target control (NTC, C4). After affinity selection, bound species were decoded by sequencing their DNA barcodes. Molecules enriched in region A, after removal of NTC (C4) signals and application of selection thresholds—were classified as putative CHI3L1 binders. **B**. Enrichment profile and representative chemical structure of the highest-ranking CHI3L1-binding small molecule identified from **DEL-C1** following affinity selection. **C**. Chemical structure of the top hit compound, **DEL-C1**.

This multi-tiered filtering pipeline effectively removed nonspecific binders and emphasized selective engagement with CHI3L1. Distinct enrichment profiles were observed across several libraries, and based on these enrichment trends (Figure 2B) and the dominant chemotypes emerging from the dataset, the top CHI3L1-binding small molecule **DEL-C1** (Figure 2C), was chosen for further evaluation. The compound was subsequently purchased from WuXi AppTec (purity > 95%) and characterized by NMR, LC-MS, mass spectrometry, HPLC purity, and chiral SFC (Figures S1-S5) to confirm its identity and suitability for downstream studies of CHI3L1 binding and functional inhibition.

### Biophysical validation

To independently confirm CHI3L1 engagement by the top DEL-identified compound, **DEL-C1**, we employed MST as an orthogonal, solution-based biophysical method. MST detects molecular interactions by monitoring fluorescence changes that occur as molecules migrate along microscopic temperature gradients. In this assay, fluorescently labeled recombinant CHI3L1 protein was titrated with increasing concentrations of **DEL-C1** under near-physiological buffer conditions. The thermophoretic signal exhibited a clear concentration-dependent shift, consistent with direct target engagement (Figure 3). Importantly, MST enables the characterization of interactions in free solution, without the need for surface immobilization, thus preserving the native conformation of CHI3L1. This is particularly advantageous for glycoproteins like CHI3L1, whose flexible and shallow binding interfaces often complicate detection using conventional binding assays. Collectively, these results validate **DEL-C1** as a CHI3L1-binding small molecule, providing orthogonal confirmation of the DEL screening hit and establishing a foundation for subsequent mechanistic and functional studies.

**Figure 3.**
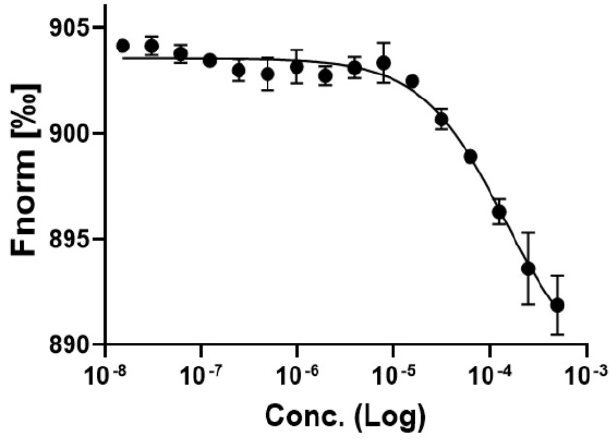
Orthogonal validation of CHI3L1 binding using MST. Dose-dependent curve for the binding of **DEL-C1** to CHI3L1 using MST (n=3).

### SPR validation

To further validate the direct interaction between CHI3L1 and the top hit compound **DEL-C1**, SPR analysis was performed under equilibrium binding conditions. Recombinant CHI3L1 protein was immobilized on a series S CM5 sensor chip, and serial dilutions of **DEL-C1** were injected in running buffer to monitor real-time association and dissociation kinetics. The resulting sensorgrams displayed a concentration-dependent increase in response units, consistent with specific and reversible binding (Figure 4). Global fitting of the response yielded an apparent dissociation constant (Kd) of 4.60 ± 2.70 µM, indicative of high-affinity interaction for small molecule binders targeting extracellular proteins. Importantly, the binding was fully reversible, with clean dissociation profiles and no evidence of nonspecific aggregation or surface interference. These data provide orthogonal confirmation of CHI3L1 engagement by **DEL-C1** and corroborate the MST findings, establishing **DEL-C1** as a validated small molecule binder suitable for downstream screening.

**Figure 4.**
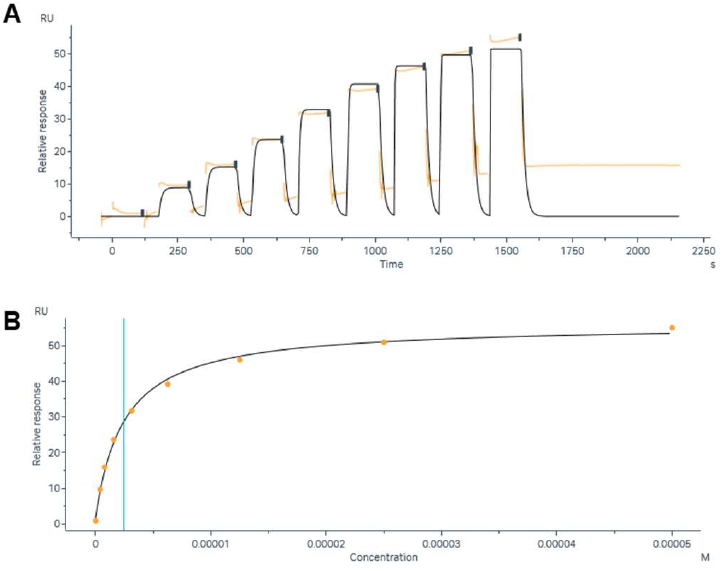
Orthogonal validation of CHI3L1 binding using SPR. **A**. Representative SPR sensorgram showing concentration-dependent binding of increasing concentrations of **DEL-C1** to immobilized recombinant CHI3L1 protein. **B**. Steady-state binding curve derived from equilibrium response values fitted to a one-site binding model, yielding an apparent dissociation constant (Kd = 4.60 ± 2.70 µM). Data are representative of three independent experiments.

### Computational analysis of the CHI3L1 binding site

Molecular docking and dynamics studies were carried out to provide insights into the binding mechanism of **DEL-C1** with CHI3L1. The docking results predicted that **DEL-C1** occupies the substrate-binding groove of CHI3L1, establishing multiple favorable interactions with key residues in the active site. The 3D and 2D binding mode analyses predicted that **DEL-C1** forms hydrogen bonds with Tyr141, Arg263 and Glu290 (Figure 5A-B). Meanwhile, **DEL-C1** was predicted to form several hydrophobic interactions with the binding site residues helping to stabilize the binding.

**Figure 5.**
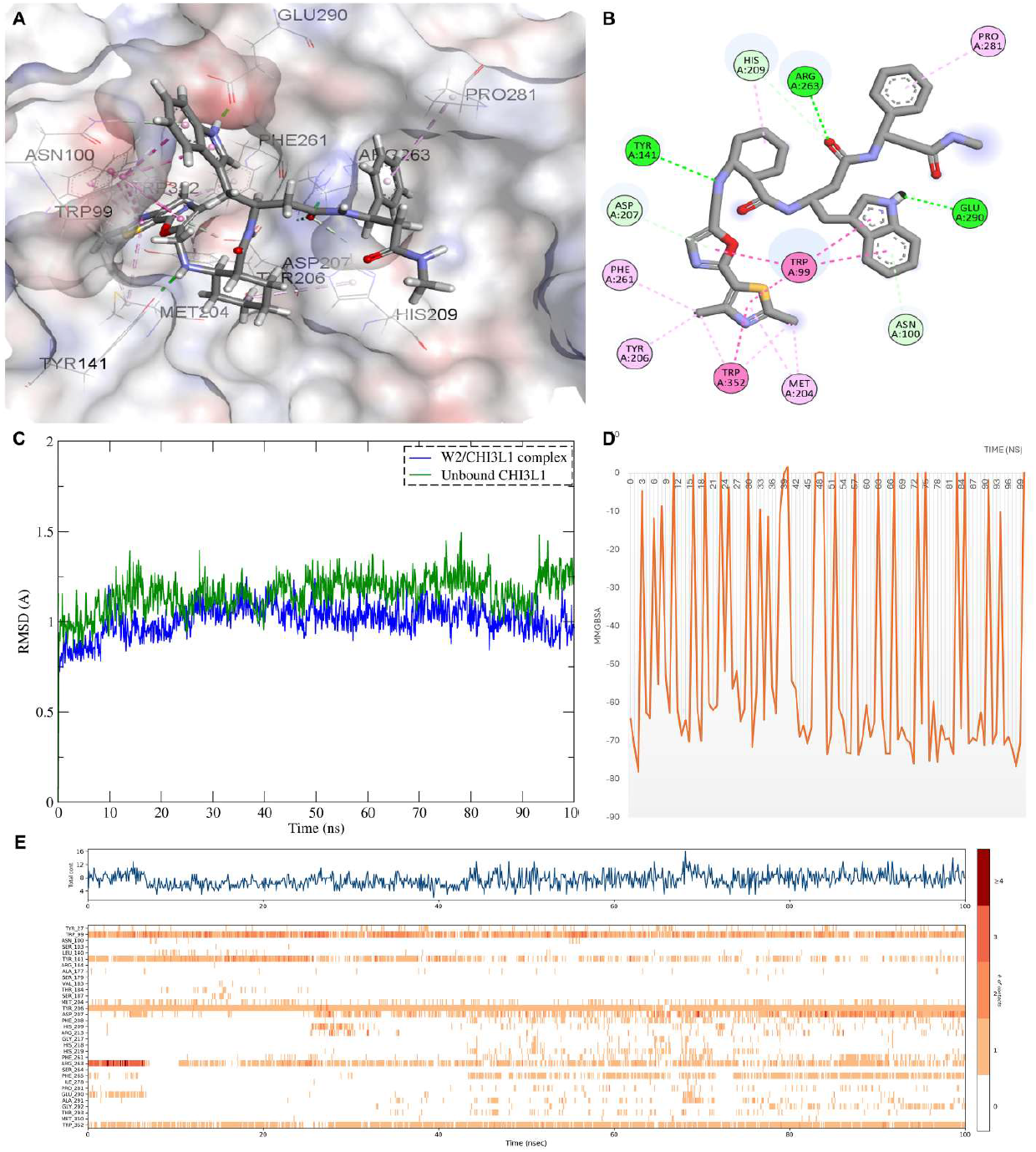
Molecular docking and dynamics simulation analysis of DEL-C1 binding to CHI3L1. **A**. Three-dimensional representation of **DEL-C1** bound to the active site of CHI3L1 (PDB ID: 8R4X, shown as surface representation). **B**. Two-dimensional interaction diagram depicting the binding mode of **DEL-C1** with CHI3L1. Green dashed lines indicate hydrogen bonds with Arg263, Tyr141, and Glu290. Pink circles represent hydrophobic interactions, green circles indicate polar residues, and purple circles show charged residues involved in ligand recognition. **C**. RMSD plot of backbone Cα atoms over 100 ns molecular dynamics simulation. **D**. MM-GBSA binding free energy plot over the simulation time. **E**. Protein-ligand contact analysis showing interaction patterns throughout the 100 ns trajectory.

The molecular dynamics simulation results supported the stability of the **DEL-C1**-CHI3L1 complex and its observed experimental binding activity. The RMSD analysis over 100 ns showed that the protein-ligand complex achieved equilibrium with an average RMSD of 1.01 Å, which is remarkably stable and indicates minimal structural perturbation upon ligand binding (Figure 5C). Notably, the RMSD of the unbound protein (average 1.15 Å) was slightly higher than the complex, suggesting that ligand binding may stabilize the protein structure by reducing conformational flexibility. Both RMSD values remained well below 2 Å throughout the simulation, confirming excellent structural stability and indicating that the docking pose represents a thermodynamically favorable binding mode.

The MM-GBSA binding free energy calculations were carried out to provide a quantitative assessment of binding affinity where the **DEL-C1**/CHI3L1 complex exhibited an average MMGBSA of -54.42 kcal/mol (Figure 5D). The predominantly negative binding energies throughout the simulation demonstrate thermodynamically favorable binding, with observed fluctuations reflecting conformational dynamics of the protein-ligand complex enhanced by the inherent flexibility of **DEL-C1** due to its multiple aliphatic linkers. The highly negative values observed at certain time points suggest transient optimization of favorable interactions, while the overall average MMGBSA indicates strong and persistent binding affinity.

The **DEL-C1**/CHI3L1 contact analysis during the 100ns MD simulation was used to predict the molecular basis for the observed stability (Figure 5E). The interaction of **DEL-C1** with Tyr206 was calculated to be key to the anchoring of **DEL-C1** within the binding site with a conserved hydrogen bond for 100% of the simulation duration. This persistent interaction suggests that Tyr206 serves as a primary anchor point for ligand recognition and binding. Additionally, **DEL-C1** formed stable π-π stacking interactions with Trp99 and Trp352 for the majority of the simulation duration which indicates that these residues contribute significantly to binding through hydrophobic and stacking interactions. The CHI3L1 binding site groove containing Tyr206 which is flanked by the two tryptophan residues (Trp99 and Trp352) appears to function as an anchoring point for the ligand and is essential for its binding.

Collectively, these findings indicate that **DEL-C1** binds to CHI3L1 with high affinity and stability through a multi-modal interaction network. The persistent engagement of Tyr206, combined with the contributions from Trp99 and Trp352, creates a robust binding interface that maintains ligand occupancy throughout the simulation. This binding mode suggests that rational optimization of **DEL-C1** could focus on enhancing interactions with these key residues to further improve binding affinity and specificity. The identification of this stable binding mode provides a structural foundation for structure-based drug design targeting CHI3L1, which has implications for therapeutic intervention in diseases where CHI3L1 plays a pathogenic role.

### Evaluation of PK and physicochemical properties of DEL-C1

The therapeutic development of small molecule modulators for AD presents unique PK challenges. Compounds must achieve sustained exposure, penetrate the blood-brain barrier (BBB), and retain safety margins suitable for chronic dosing in elderly patients. To evaluate the drug-likeness of **DEL-C1**, we conducted a comprehensive in vitro ADME assessment (Table 1).

**Table 1.**
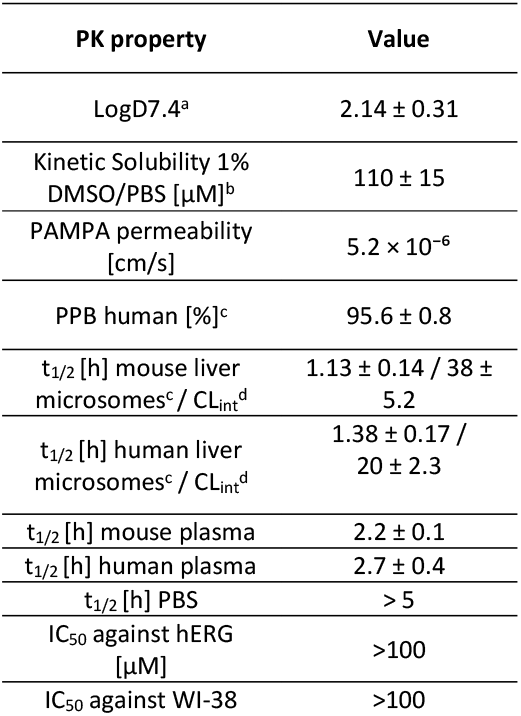
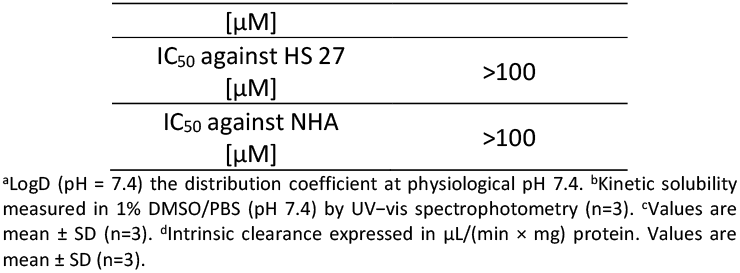
In vitro PK profile of DEL-C1.

**DEL-C1** exhibited a LogD_7_._4_ of 2.14 ± 0.31, reflecting a balanced hydrophilic–lipophilic profile favorable for oral bioavailability and BBB penetration. Compounds within the 1.5-3.0 LogD window typically display optimal CNS permeability while minimizing efflux via P-glycoprotein transporters. The kinetic solubility of 110 ± 15 µM further supports developability, suggesting that the compound can achieve adequate free-drug concentrations without formulation enhancement. Its PAMPA permeability (5.2 × 10^−6^ cm s^−1^) places it in the high-permeability class, consistent with passive diffusion across the BBB and supporting central exposure under systemic administration.In metabolic assays, **DEL-C1** demonstrated moderate intrinsic clearance (Table 1), with t_1/2_ values of 1.13 h (mouse) and 1.38 h (human) corresponding to CL_int_ values of 38 ± 5.2 and 20 ± 2.3 µL min^−1^ mg^−1^, respectively. These rates suggest the compound is metabolically stable enough to maintain therapeutic levels yet sufficiently cleared to avoid accumulation during repeated dosing. The lower clearance in human microsomes indicates improved metabolic resilience relative to mouse, which is desirable for translational predictability. Plasma stability studies (Table 1) revealed half-lives of 2.2 ± 0.1 h (mouse) and 2.7 ± 0.4 h (human), consistent with limited degradation by plasma esterases or oxidative pathways, while PBS stability exceeding 5 h confirms chemical integrity in physiologic buffer. These parameters indicate that metabolic processes, not chemical instability, govern systemic clearance. Plasma protein binding of 95.6 ± 0.8 % ensures a pharmacologically active free fraction while providing a depot effect that extends circulation time. Such moderate binding is typical of CNS drugs, balancing target engagement and tissue distribution.

From a safety perspective, **DEL-C1** exhibited no measurable hERG channel inhibition (IC_50_ > 100 µM), minimizing concerns for cardiotoxicity. Cytotoxicity assays across non-cancerous human cell lines (WI-38 fibroblasts, HS-27 stromal cells, and normal human astrocytes) showed IC_50_ values above 100 µM, confirming an excellent safety margin and low off-target reactivity. The absence of astrocytic toxicity is particularly important for a compound designed to modulate an astrocyte-derived target such as CHI3L1. Collectively, these data support the advancement of **DEL-C1** as a lead CHI3L1 modulator for AD, combining chemical tractability with a PK profile conducive to CNS exposure. The balanced clearance and plasma stability of **DEL-C1** suggest a PK half-life compatible with once- or twice-daily oral dosing, while its physicochemical parameters predict brain/plasma ratios favorable for engaging astrocytic CHI3L1. Future in vivo studies will quantify brain penetration, confirm exposure-response relationships, and evaluate the ability of **DEL-C1** to attenuate neuroinflammatory and neurogenic deficits in transgenic AD models.

### Functional validation as a potential AD drug candidate

To further establish the role of CHI3L1 in astrocyte-driven neuroinflammation, we investigated whether pharmacological inhibition of CHI3L1 could restore astrocytic homeostasis under inflammatory stress. CHI3L1 upregulation in reactive astrocytes has been linked to IL-6 secretion, lysosomal alkalinization, and impaired Aβ uptake in AD. Given this, we evaluated whether **DEL-C1** could reverse these pathological phenotypes. Notably, astrocytes play a central role in Aβ clearance, and dysfunction in astrocytic endolysosomal trafficking contributes to Aβ accumulation in AD. Recent studies have shown that CHI3L1 overexpression in reactive astrocytes disrupts lysosomal acidification and phagocytic capacity via CRTH2- and RAGE-mediated signaling.^14–18^

As shown in Figure 6A, recombinant CHI3L1 (300 ng mL^−1^) reduced the uptake of pHrodo-Aβ_1−42_ to 70 ± 8 % of vehicle, consistent with a loss of endocytic activity. Treatment with **DEL-C1** restored uptake in a dose-dependent manner, reaching baseline levels at 25 µM. Lysosomal degradation of DQ-BSA (Dye-Quenched Bovine Serum Albumin) was similarly improved, from 0.66 ± 0.07 fold to 0.97 ± 0.06 fold (Figure 6B) and the elevated LysoSensor ratio (1.22 ± 0.07) induced by CHI3L1 normalized to 1.01 ± 0.06, indicating restored acidification (Figure 6C). **DEL-C1** also attenuated CHI3L1-driven cytokine release, reducing secreted CHI3L1 and IL-6 without affecting cell viability (Figures 6D,E). These findings indicate that **DEL-C1** reverses CHI3L1-induced astrocyte dysfunction by restoring lysosomal proteolysis and inflammatory balance. By targeting CHI3L1, **DEL-C1** disrupts the pathological CHI3L1-CRTH2/RAGE axis, preventing IKKβ-S6K1-S6 activation and preserving vesicular pH homeostasis. This functional rescue parallels that observed in CHI3L1-knockout astrocytes, which exhibit enhanced Aβ phagocytosis and reduced IL-6 production. Collectively, the results demonstrate that **DEL-C1** acts as a dual-function modulator that suppresses CHI3L1-driven neuroinflammation while restoring astrocyte-mediated Aβ clearance, two interrelated processes central to AD progression.

**Figure 6.**
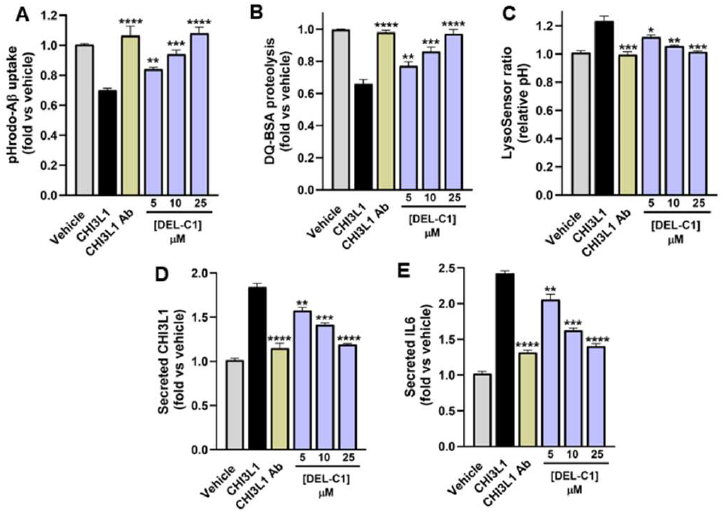
DEL-C1 restores astrocytic Aβ clearance and lysosomal function disrupted by CHI3L1. **A**. Flow-cytometric quantification of pHrodo-Aβ_1−42_ uptake in human iPSC-derived astrocytes shows that recombinant CHI3L1 (300 ng mL^−1^) markedly reduced Aβ uptake to 70 ± 8 % of vehicle, whereas **DEL-C1** restored uptake in a dose-dependent manner (5-25 µM). **B-C**. Lysosomal proteolysis, measured by DQ-BSA degradation, and lysosomal pH (LysoSensor ratio) confirmed recovery of degradative capacity and acidification following **DEL-C1** or α-CHI3L1 antibody treatment (10 µg mL^−1^). **D-E. DEL-C1** attenuated CHI3L1-driven cytokine release, reducing secreted CHI3L1 and IL-6 without affecting cell viability. Data represent mean ± SD from n = 3 independent experiments performed in triplicate. Statistical significance was determined by one-way ANOVA with Dunnett’s post hoc test versus the CHI3L1 group: *p* < 0.05 (*), *p* < 0.01 (**), *p* < 0.001 (***), and *p* < 0.0001 (****).

To confirm that **DEL-C1** directly modulates the inflammatory signaling cascade triggered by CHI3L1, we employed a NF-κB-luciferase reporter assay in human iPSC-derived astrocytes. CHI3L1 is known to activate NF-κB transcription through CRTH2-dependent IKKβ phosphorylation, leading to cytokine release and sustained glial activation. As shown in Figure 7A, stimulation with recombinant CHI3L1 (300 ng mL^−1^, 6 h) increased NF-κB-driven luciferase activity to 2.6 ± 0.3 fold relative to vehicle. Preincubation with **DEL-C1** resulted in a concentration-dependent suppression of NF-κB activation, reducing signal to 1.9 ± 0.2 fold at 5 µM and 1.3 ± 0.1 fold at 25 µM. A neutralizing anti-CHI3L1 antibody (10 µg mL^−1^) produced a comparable reduction (1.2 ± 0.1 fold). No change in Renilla normalization or cell viability (Figure 7B) was observed, confirming that signal reduction reflected pathway inhibition rather than cytotoxicity.

**Figure 7.**
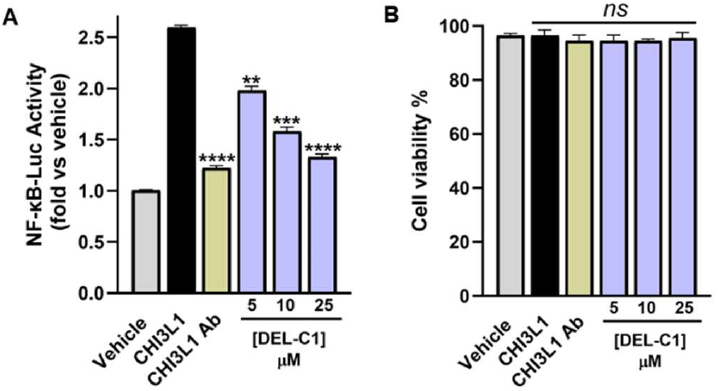
DEL-C1 inhibits CHI3L1-induced NF-κB activation in human astrocytes. **A**. NF-κB-luciferase reporter assay showing that recombinant CHI3L1 (300 ng mL^−1^, 6 h) strongly activates NF-κB transcription in human iPSC-derived astrocytes. Pretreatment with **DEL-C1** (5-25 µM) suppressed NF-κB activity in a concentration-dependent manner, restoring levels to near baseline at 25 µM. The neutralizing anti-CHI3L1 antibody (10 µg mL^−1^) produced a similar reduction in NF-κB activity, confirming pathway specificity. **B**. Cell viability remained unaffected across all conditions, indicating that signal inhibition was not due to cytotoxicity. Data represent mean ± SD from n = 3 independent experiments performed in triplicate. Statistical significance was determined by one-way ANOVA with Dunnett’s post hoc test versus the CHI3L1 group: *ns* non-significant, *p* < 0.01 (**), *p* < 0.001 (***), and *p* < 0.0001 (****).

These results establish that **DEL-C1** effectively blocks CHI3L1-induced NF-κB transcriptional activation, providing a molecular correlate to its functional rescue of lysosomal integrity and cytokine balance. The combined data demonstrate that **DEL-C1** interrupts the core inflammatory loop downstream of CHI3L1 engagement, offering both mechanistic insight and translational relevance for CHI3L1-targeted modulation of astrocytic neuroinflammation in AD.

## Conclusion

This work presents the discovery and validation of a small molecule directly targeting CHI3L1, a key astrocytic driver of neuroinflammation in AD. Through a combination of DEL screening, biophysical validation, molecular modeling, and cellular assays, we identified **DEL-C1** as a CHI3L1 binder with favorable CNS drug-like properties. **DEL-C1** demonstrated consistent binding across MST and SPR, exhibited a stable interaction network with key residues in the CHI3L1 groove, and showed a balanced PK profile supportive of brain penetration. Importantly, **DEL-C1** restored Aβ uptake and lysosomal function while attenuating CHI3L1-driven IL-6 secretion and NF-κB activation, effectively reprogramming astrocyte reactivity toward a homeostatic state. These findings provide the first experimental evidence that CHI3L1 is a tractable, druggable target amenable to small molecule modulation. **DEL-C1** thus represents both a mechanistic probe and a promising lead compound for therapeutic strategies aimed at mitigating neuroinflammation and neurodegeneration in AD.

## Experimental

### DEL Screening

A DNA-encoded chemical library (DEL) comprising 4.2 billion compounds, supplied by WuXi AppTec, was utilized to screen for interactions with the human CHI3L1 protein. Prior to conducting affinity selection, a protein capture assay was performed to confirm the stable immobilization of CHI3L1 protein (Cat. #CH1-H5228, ACROBiosystems, Newark, DE, USA) onto the affinity matrix. This preliminary validation led to the optimization of the selection buffer, which consisted of 1x PBS, 0.05% Tween-20, 0.1 mg/mL sssDNA, and 10 mM imidazole. The elution buffer was formulated with 1x PBS and 0.05% Tween-20. Each round of selection employed 5 µg of CHI3L1 protein in conjunction with HisPur™ Ni-NTA Magnetic Beads (Cat. # 88831, Thermo Fisher, Waltham, MA, USA) as the affinity matrix.

Affinity selection, encompassing both protein immobilization and subsequent selection rounds, was executed in accordance with the manufacturer’s protocol. The DELopen kit included four identical sets of 27 distinct libraries, collectively covering 4.2 billion compounds. One set was allocated for CHI3L1 screening, while two others were used for screening different human proteins (undisclosed). The fourth set functioned as a non-target control (NTC), using HisPur™ Ni-NTA Magnetic Beads without protein to assess nonspecific binding.

Following the second selection round, the heated elution sample underwent a self-QC step, ensuring the concentration of selected DEL molecules remained below 5×10_8_ copies (standard control) and free from cross-contamination. This post-selection sample was submitted to WuXi AppTec, where it passed sequencing quality control before advancing to next-generation sequencing (NGS) for decoding the DNA-linked small molecules. Data analysis revealed the top-binding compound. WuXi AppTec disclosed the structure to the authors, and upon request, synthesized tag-free version for subsequent investigation.

### MST assay

MST was conducted on a Monolith NT.115 (NanoTemper Technologies) to assess binding between hit compound and recombinant His-tagged human CHI3L1 (Cat. #CH1-H5228, Acro Biosystems). The protein was fluorescently labeled with the Monolith His-Tag Labeling Kit RED-tris-NTA, 2nd Generation (Cat. #MO-L018) and diluted to 80 nM. Test compounds were prepared as a 16-point serial dilution (500 µM to low nM) in assay buffer (10 mM HEPES, 150 mM NaCl, 0.1% Pluronic F-127, 1 mM TCEP, 8% DMSO, pH 7.4). Protein and compound solutions were mixed (1:1) maintaining a final DMSO concentration of 4%, incubated for 30 min at room temperature, centrifuged briefly, and loaded into capillaries. Measurements were taken using standard LED and IR-laser settings.

### SPR assay

The SPR validation of CHI3L1 binding was performed using Biacore™ 8K SPR system (Cytiva, Marlborough, MA, USA) following our previously reported method.^23,25^ hCHI3L1-His Protein (8.0 μg/mL in acetate buffer, pH 5.0) was immobilized on a Series S Sensor Chip CM5 (29104988, Cytiva, Marlborough, MA, USA) using a commercial amine coupling kit (BR100050, Cytiva, Marlborough, MA, USA). Immobilization was performed at a flow rate of 10 μL/min for 420 s, reaching an immobilization level around 3000 RU, followed by blocking with ethanolamine. The flow cell with solely ethanolamine-block on the same channel served as the reference. Immobilization buffer: PBS-P+ (28995084, Cytiva, Marlborough, MA, USA). Briefly, gradient concentrations of the compound were prepared in the assay buffer, and injected over the sensor chip in a single-cycle kinetics model with a flow rate at 30 μL/min for 120 s per injection. After each injection, a 30 s-regeneration was performed at a flow rate of 30 μL/min using the regeneration buffer. Data were analyzed using Biacore™ Insight Evaluation Software (Cytiva, Marlborough, MA, USA). The assays were performed in a single time experiment and the Kd values were given as the exact numberm or in individual triplicate and the Kd values were given as mean ± SD.

### Computational study

Molecular docking simulations were performed using Schrödinger Suite 2023.4. The crystal structure of CHI3L1 was retrieved from the Protein Data Bank (PDB ID: 8R4X^30^). The protein structure was prepared using Maestro’s protein preparation wizard at default settings which included addition of hydrogen atoms, assignment of appropriate protonation states at pH 7.0, optimization of hydrogen bond networks, and energy minimization using the OPLS4 force field. The ligand **DEL-C1** was prepared using LigPrep module with default parameters, generating possible ionization states at pH 7.4 ± 2.0.

Receptor grid generation was performed centered on the active site binding pocket using the native co-crystal (XZ0, a validated CHI3L1 inhibitor also called CHI3L1-IN-1). Molecular docking was carried out using Glide module in standard precision (SP) mode. Visualization of the binding mode and the 2D interaction diagram were generated to identify key residues involved in ligand-protein interactions. Subsequently, two independent 100ns MD simulations were carried out using the DESMOND software according to our previously published methodology^23^ to compare the **DEL-C1**/CHI3L1 complex with the unbound protein.

### Evaluation of PK and physicochemical properties

The These experiments were conducted following our previously reported methods.^25^ The evaluations included LogD7.4 determination, microsomal stability, kinetic solubility, and cytotoxicity profiling across multiple cell lines. Solubility was assessed using UV–visible spectrophotometry, while cell viability was measured using the PrestoBlue assay.

### Cell culture and reagents

Human iPSC-derived astrocytes were maintained according to the manufacturer’s protocol in astrocyte medium (DMEM/F12 supplemented with 10% FBS) at 37 °C in a humidified atmosphere of 5% CO_2_. Recombinant human CHI3L1 (YKL-40) was purchased from R&D Systems. For comparison and pathway validation, a neutralizing anti-CHI3L1 antibody (R&D Systems) was used as a positive control. All assays were performed in 96-well format unless otherwise indicated.

### Astrocyte Aβ Uptake and Lysosomal Function Assay

Human iPSC-astrocytes were seeded at 2 × 10^4^ cells per well (96-well plates) and cultured for 48 h. Cells were pre-treated with **DEL-C1** (5-25 µM, 30 min) prior to stimulation with CHI3L1 (300 ng mL^−1^, 24 h). For Aβ uptake, cells were incubated with pHrodo™-Red– labeled Aβ_1−42_ (1 µM) for 2 h, dissociated using Accutase, fixed in 2% paraformaldehyde, and analyzed on a BD Fortessa flow cytometer (Ex/Em 560/585 nm). The mean fluorescence intensity (MFI) from ≥10,000 live events per well was normalized to vehicle controls.

Lysosomal degradation was assessed using DQ-BSA Green (10 µg mL^−1^, 2 h), and lysosomal pH was measured using LysoSensor™ DND-160 (Ex/Em 340/440–535 nm ratio). Supernatants were collected for CHI3L1 and IL-6 quantification by ELISA (R&D Systems). Cell viability was assessed using CellTiter-Glo® (Promega). All measurements were performed in triplicate from at least three independent experiments.

### NF-κB Reporter Assay

To evaluate the effect of **DEL-C1** on CHI3L1-driven inflammatory signaling, human iPSC-astrocytes were transduced with an NF-κB-firefly luciferase reporter and a Renilla luciferase internal control. Cells were maintained for 72 h post-transduction to achieve stable expression before treatment. Cells were pre-incubated with **DEL-C1** (5-25 µM, 30 min) followed by stimulation with CHI3L1 (300 ng mL^−1^, 6 h). Firefly and Renilla luciferase activities were measured using the Dual-Glo® Luciferase Assay System (Promega). Results were expressed as normalized Firefly/Renilla ratios and presented as fold-change relative to untreated vehicle controls. Cell viability was verified in parallel using CellTiter-Glo®.

## Acknowledgments

This work was supported by the National Institute of Neurological Disorders and Stroke under grant number R01NS136524 (PI: Gabr).

