## Supporting Information for "Discovery of a CHI3L1-Targeted Small Molecule Modulating Neuroinflammation in Alzheimer’s Disease via DNA-Encoded Library (DEL) Screening^†^"

*Electronic Supplementary Information*

| **Contents** |  |
| --- | --- |
| ^1^H NMR spectrum of compound **DEL-C1** in DMSO-*d*6 | S2 |
| LCMS report of compound **DEL-C1** | S3 |
| Mass spectrum of compound **DEL-C1** | S4 |
| HPLC purity of compound **DEL-C1** | S5 |
| Chiral SFC report of compound **DEL-C1** | S6 |

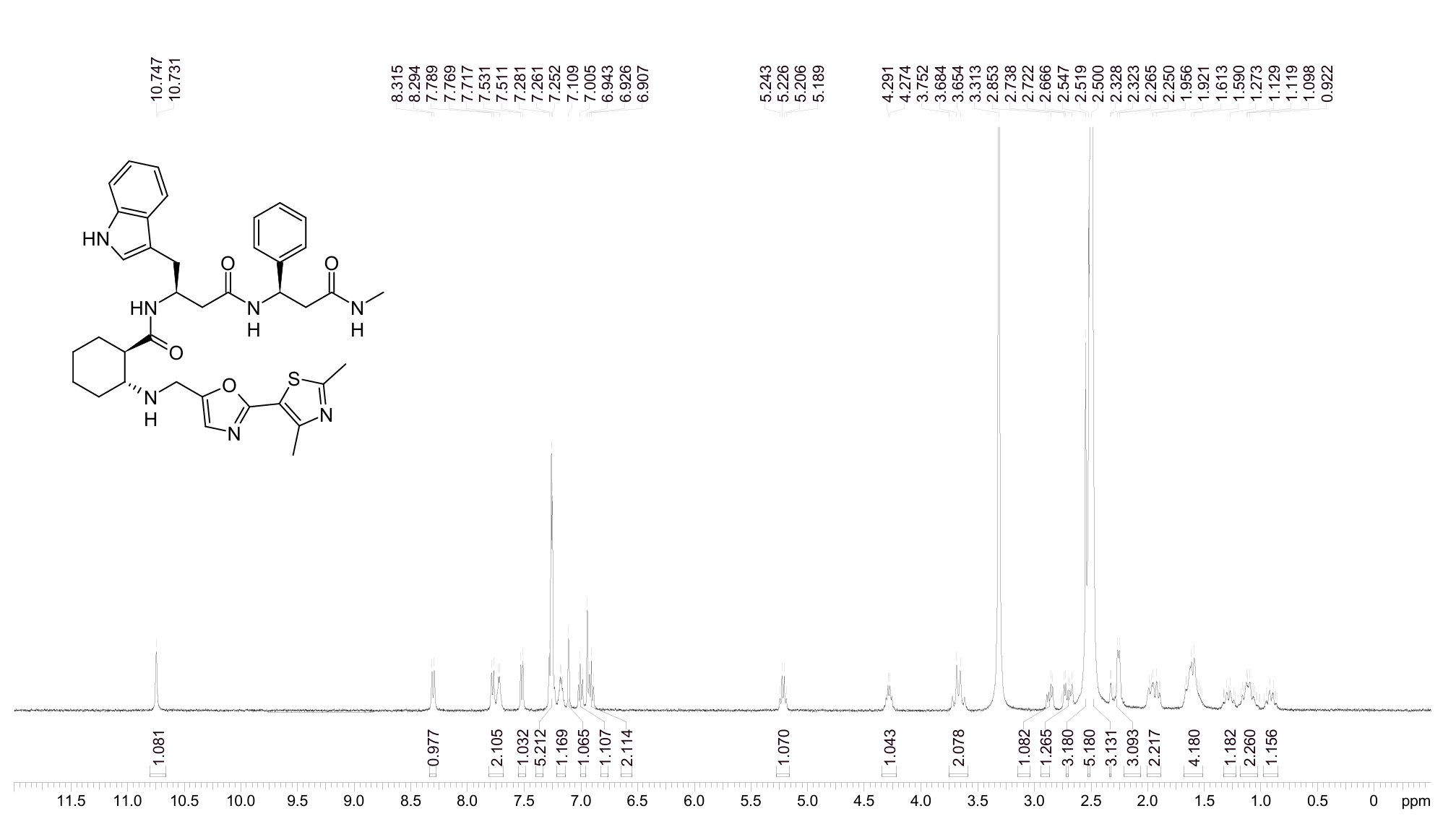

**Figure S1**. ^1^H NMR spectrum of compound **DEL-C1** in DMSO-*d*6.

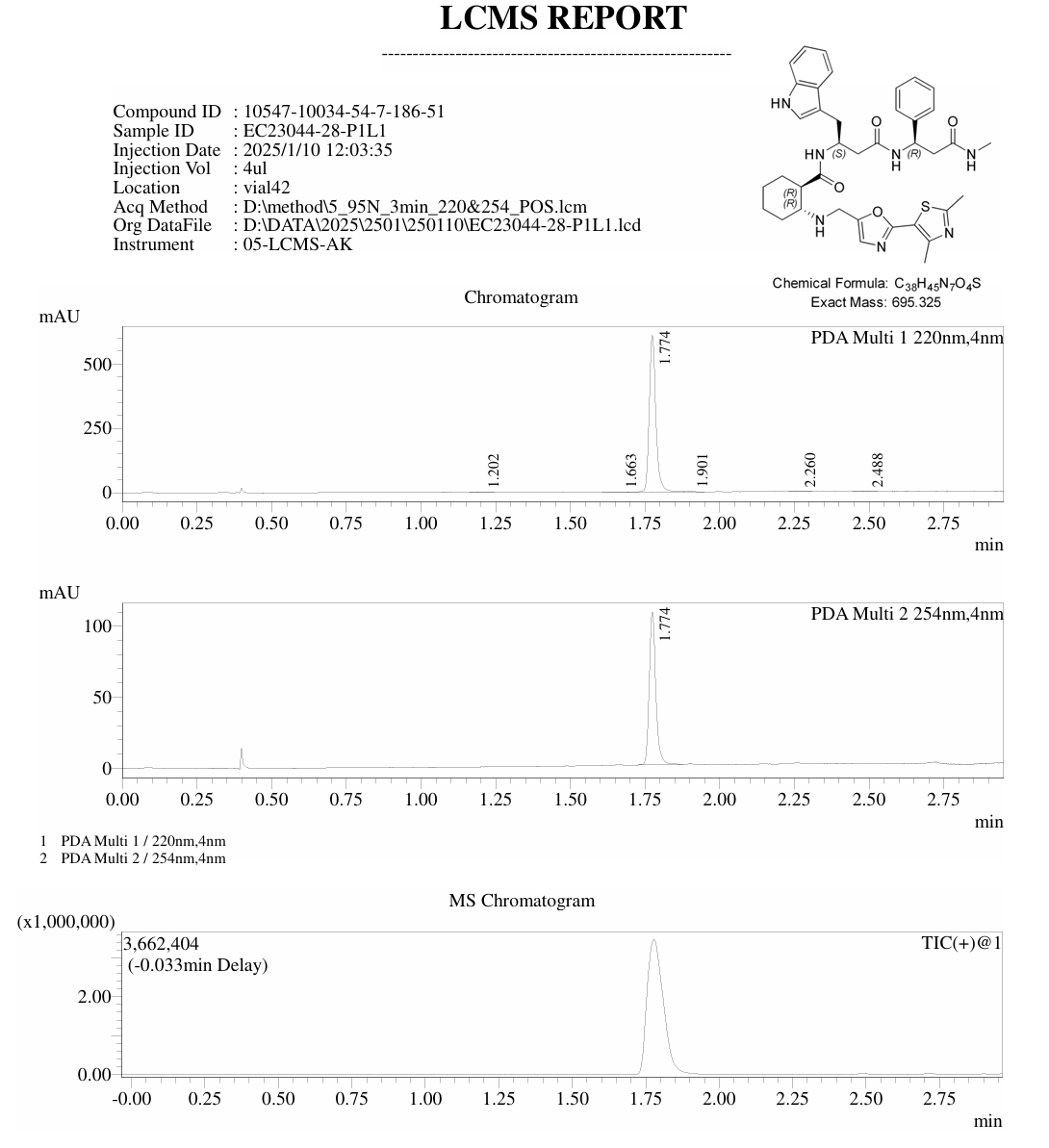

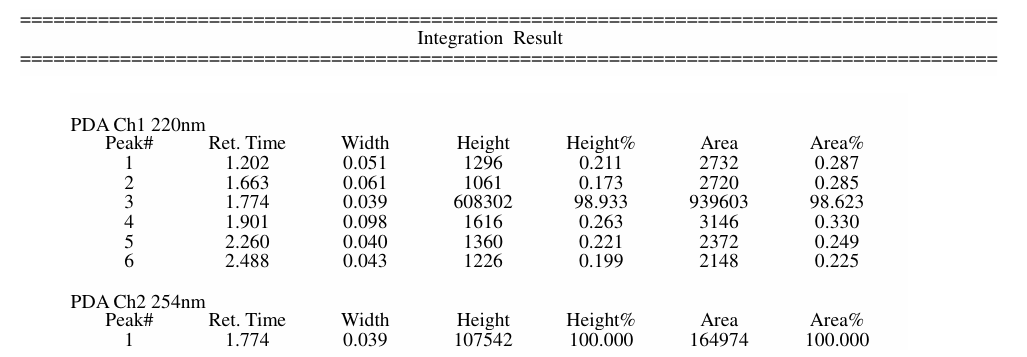

**Figure S2**. LCMS report of compound **DEL-C1**.

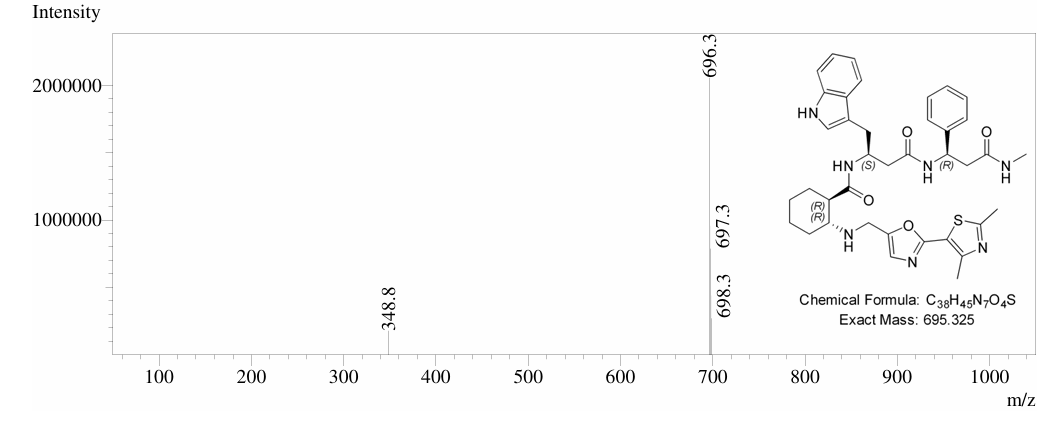

**Figure S3**. Mass spectrum of compound **DEL-C1**.

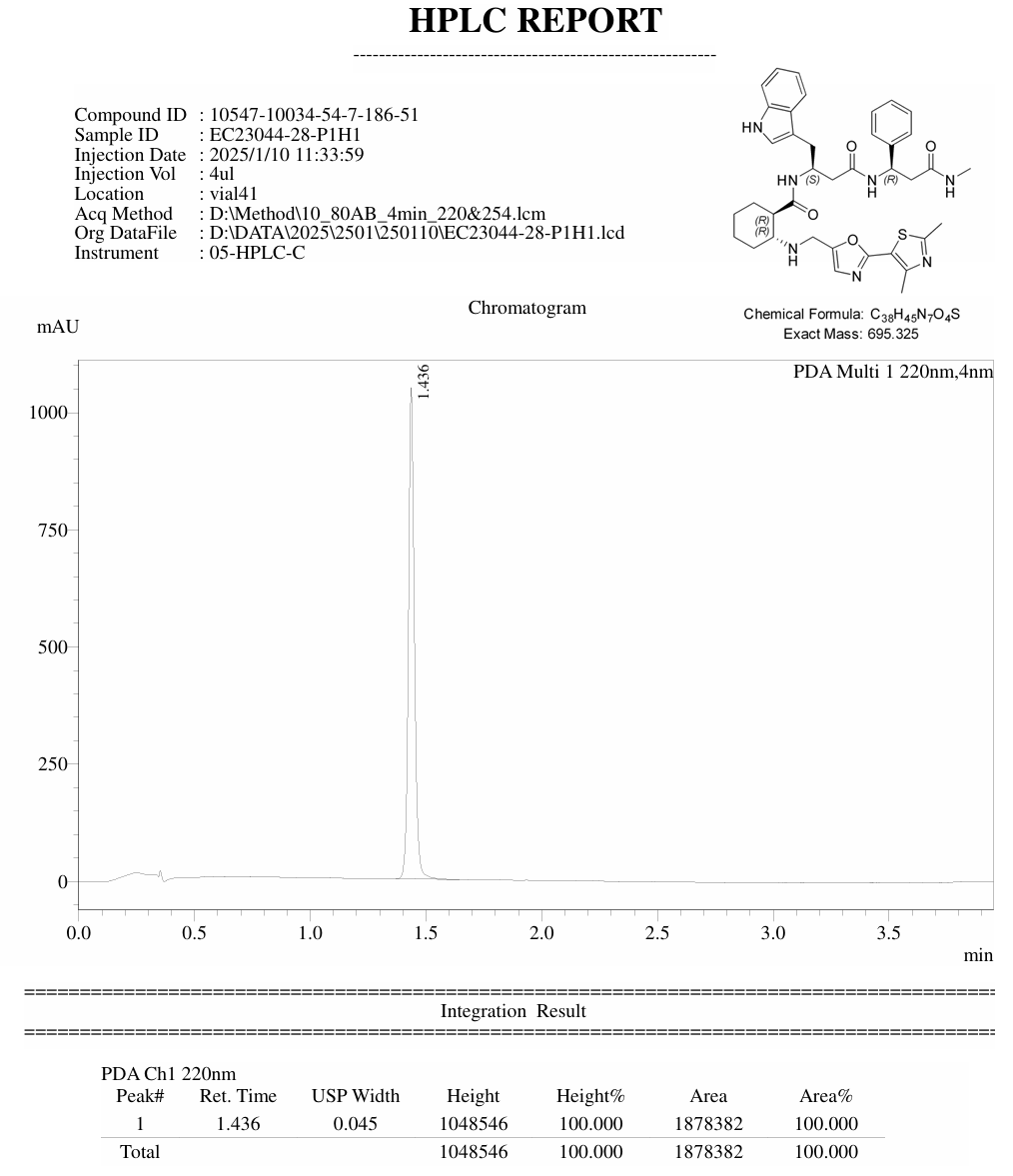

**Figure S4**. HPLC purity of compound **DEL-C1**.

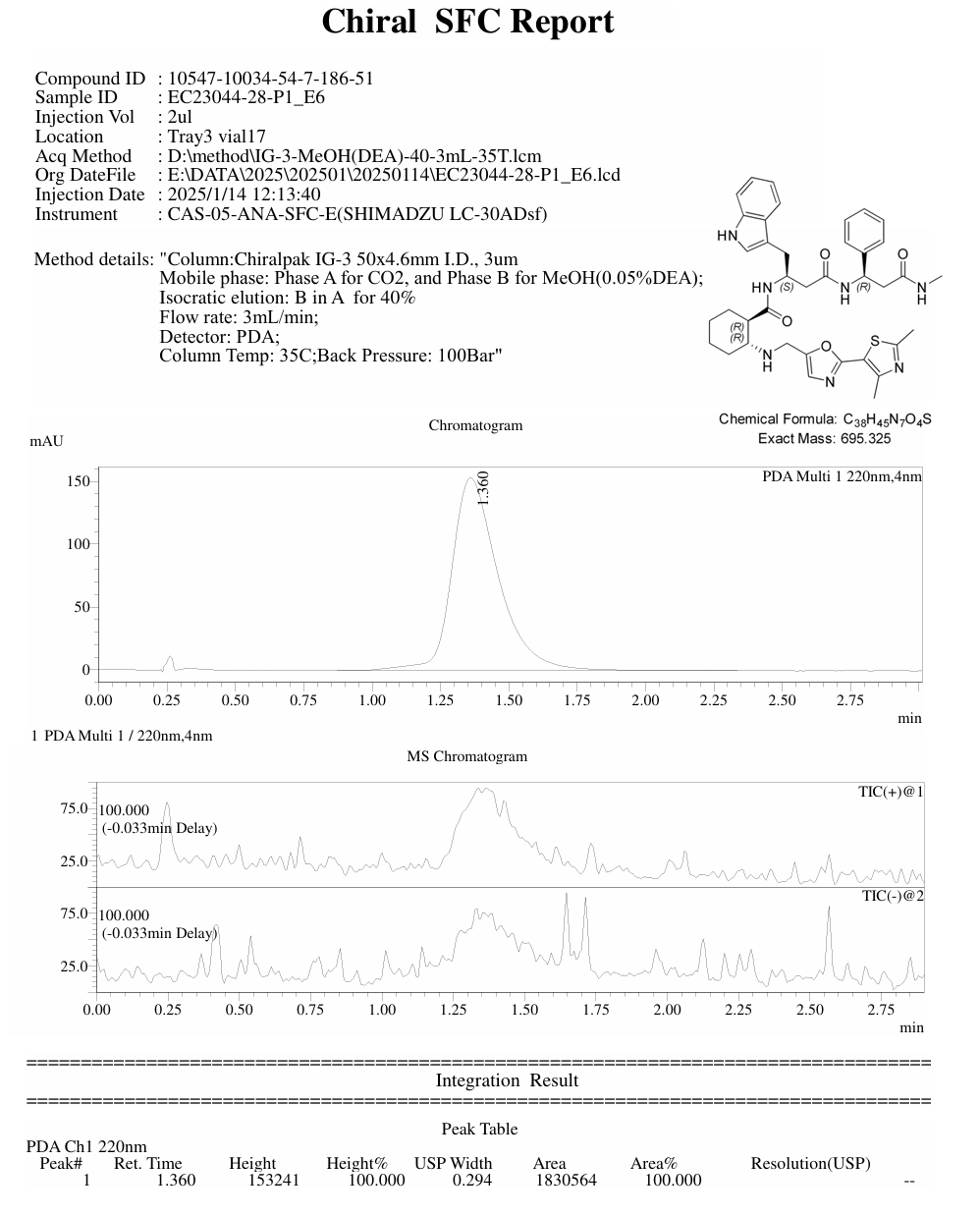

**Figure S5**. Chiral SFC report of compound **DEL-C1**.
